# funspace: an R package to build, analyze and plot functional trait spaces

**DOI:** 10.1101/2023.03.17.533069

**Authors:** Carlos P. Carmona, Nicola Pavanetto, Giacomo Puglielli

## Abstract

1. Functional trait space analyses are pivotal to define species’ ecological strategies across the tree of life. Yet, there is no single application that streamlines the many sometimes-troublesome steps needed to build and analyze functional trait spaces.
2. To fill this gap, we propose `funspace`, an R package to easily handle bivariate and multivariate (PCA-based) functional trait space analyses. The six functions that constitute the package can be grouped in three modules: ‘Building and exploring’, ‘Mapping’, and ‘Plotting’.
3. The building and exploring module defines the main features of a functional trait space (e.g., functional diversity metrics) by leveraging kernel density-based methods. The mapping module uses general additive models to map how a target variable distributes within a trait space. The plotting module provides many options for creating flexible and high-quality figures representing the outputs obtained from previous modules. We provide a worked example to demonstrate a complete `funspace` workflow.
4. `funspace` will provide researchers working with functional traits across the tree of life with an indispensable asset to easily explore: (i) the main features of any functional trait space, (ii) the relationship between a functional trait space and any other biological or non-biological factor that might contribute to shaping species’ functional diversity.

## Introduction

Traits are defined as characteristics of individual organisms that can be measured at any relevant level of biological organization (Dawson et al., 2021). They reflect species adaptations to the main selective forces in their natural habitats, and therefore have a (more or less direct) link with species’ performance - i.e., functional traits - and/or ecosystem processes (de Bello et al., 2021). Thus, functional traits represent an essential tool for ecologists and ecophysiologists worldwide to describe the diversity of organisms’ form and function in relation to any aspect that might influence a species’ biology.

Classical studies have hypothesized that species adaptations to the environment might have evolved around few theoretical strategic axes (Greenslade, 1983; Grime, 1977; Pianka, 1970; Westoby, 1998). Subsequent research has attempted to characterize such axes using functional traits for fish (Winemiller & Rose, 1992), corals (Darling et al., 2012), and vascular plants (e.g., Díaz et al., 2004), among other groups of organisms. The implication of having a handful of axes describing species’ strategies implies that the diversity of organisms’ form and function must be constrained by trade-offs among traits (Pianka, 1970 for animals; Greenslade, 1983 for insects; Westoby et al., 2002 for vascular plants). Nowadays, we know that pervasive trade-offs between traits constrain organisms’ form and function along (relatively) few trait dimensions that are often independent, thus limiting possible trait combinations within two- or highly dimensional spaces, also called functional trait spaces, across the tree of life (Carmona, Bueno, et al., 2021; Carmona, Tamme, et al., 2021; Díaz et al., 2016; Junker et al., 2023; Mouillot et al., 2021; Westoby et al., 2021; Winemiller et al., 2015). Expressing the diversity of organisms’ form and function within functional trait spaces of a reduced number of dimensions allows for robust generalization and quantification of species’ strategic axes of trait variation (see Mouillot et al., 2021). Thus, functional trait spaces represent indispensable tools to describe organisms’ functional diversity across the tree of life.

The most common quantitative tool for building multivariate functional trait spaces is Principal Component Analysis (PCA), as it allows reducing a trait dataset to few independent trait dimensions defined by the inherent relationships between traits. Recent methodological advances have permitted to explore additional features of functional trait spaces besides the trait dimensions that define them. In particular, kernel density analyses (Carmona et al., 2016, 2019; Duong, 2007) have revealed that species occupy functional trait spaces differentially, resulting in species clumping around some trait combinations that are much more frequent than others (Carmona, Bueno, et al., 2021; Díaz et al., 2016; Puglielli et al., 2021 for vascular plants; Carmona, Tamme, et al., 2021; Cox et al., 2021; Toussaint et al., 2021 for different animal groups). As a result, kernel density estimates are nowadays widely employed to estimate different properties about the way in which different species and groups of species occupy a functional trait space, including aspects such as the amount of space occupied (functional richness) or the degree to which species in a group have different trait combinations (functional divergence), among other indices of functional diversity (see Mammola et al., 2021 for a review).

A first pitfall arising when using PCA is that missing values are not allowed in the dataset. This is especially the case of large-scale analyses, where the unprecedented trait availability provided by global databases contrasts with the use of disparate trait frameworks across studies and disciplines, resulting in large databases including many traits but with missing information for many of the species (e.g., the TRY plant trait database, Kattge et al., 2020). This has pushed research towards developing imputation methods (e.g., Penone et al., 2014; Schrodt et al., 2015; Stekhoven & Bühlmann, 2012) that mostly use machine-learning techniques to impute missing trait values based on trait-trait correlations sometimes improved by accounting for species’ phylogenetic information (Penone et al., 2014). A key point to consider is that missing trait information is not distributed randomly within datasets. For example, across mammals, big species with large range areas are generally better informed than small species or species with small ranges (González-Suárez et al., 2012); similar biases have been described for plants (Carmona, Bueno, et al., 2021; Sandel et al., 2015). Recent research is increasingly suggesting that imputation can largely correct these biases, so that functional diversity patterns inferred from imputed datasets are much closer to the real ones than those that would be estimated using only species with complete trait information (Penone et al., 2014; Stewart et al., 2022).

Even when there are no missing data, another pitfall is deciding how many dimensions to retain to maximize the information contained in the dataset while minimizing information redundancy. Usually, this is done by retaining the first two Principal Components (PCs) since they capture most of the variance in a dataset. However, if the intrinsic dimensionality of the considered trait dataset is higher, this approach might lead to the loss of biologically relevant information (Laughlin, 2014). Several methods have been proposed to determine the number dimensions (Mouillot et al., 2021; Peres-Neto et al., 2005). In the context of PCA, the parallel analysis method proposed by Horn, 1965 is one of the most precise methods in identifying the correct number of principal components to retain (Peres-Neto et al., 2005), and one of the most widely used. This method contrasts eigenvalues produced through a PCA on *n* random data sets of uncorrelated variables with the same dimension of the original dataset to produce eigenvalues for components that are adjusted for the sample error-induced inflation. Apart from some large cross-species studies (e.g., Carmona, Bueno, et al., 2021; Carmona, Tamme, et al., 2021; Díaz et al., 2016; Guillemot et al., 2022; Toussaint et al., 2021), the two described pitfalls of using PCA are hardly considered when building functional trait spaces. We argue that this might mostly depend on lack of knowledge of the existence of specific procedures, and likely, on the potential difficulties when implementing them. However, the most common practices when building functional trait spaces remain choosing the first two PCs and/or dropping observations with missing values, with the inevitable loss of biological information.

Another aspect that deserves attention is how to use functional trait spaces as a ground to test for the relationship between species’ trait strategies and additional variables. This includes analyses such as exploring the relationship between species’ ecological strategies (i.e., their position in the functional trait space) and climatic features at a species’ habitat, or to explore how extinction risk (Carmona, Tamme, et al., 2021), as well as any other adaptive syndrome (Puglielli et al., 2022), is related to species traits. However, analyzing these multidimensional relationships is inherently complex (Villéger et al., 2011). One layer of complexity is provided by the need to use the multiple dimensions defining the functional trait space as predictors in models attempting to link multidimensional species strategies to other ecological dimensions (e.g., climate). Another problem is that more than often we do not have any previous knowledge on the functional form (e.g., linearity) of the relationship linking the target dimension to species’ strategies. In addition, visualizing such relationships is not straightforward, and we often rely on separately analyzing relationships between response variables and single trait dimensions (e.g., a single PC). These difficulties can be overcome considering functional trait spaces as a set of coordinates that we can use to build maps of any response variable of interest. General additive models (GAMs), used to estimate smooth functional relationships between predictor variables and a response (Pedersen et al., 2019), provide a solution to do that. Accordingly, they have been consistently used for fitting and mapping, for instance, species distribution models’ predictions in spaces defined by geographical coordinates (e.g., Naimi & Araújo, 2016). GAMs allow expressing the models’ predictor as a multidimensional smoother, allowing to use the functional trait space dimensions as a single predictor (see Carmona, Tamme, et al., 2021; Puglielli et al., 2022; for examples). In the case of functional trait spaces, GAMs provide a sound and flexible modeling solution because they operate using piecewise functions adapting to the local conditions within the functional trait space, so that GAMs behavior in a particular portion of the data point cloud does not overall alter the global behavior of the model (Pedersen et al., 2019; Venables & Dichmont, 2004). While this comes at the cost of the interpretability of the coefficients, spline regressions are more easily interpreted from a graphical point of view rather than through the values of their coefficients (Venables & Dichmont, 2004). Thus, GAMs become particularly useful to visualize and test patterns of how target variables vary within functional trait spaces.

Despite functional trait spaces (bivariate and multivariate) are widely used in disparate research fields spanning ecology, plant science, animal ecology, evolutionary ecology – e.g., searching the term “trait space” in Web of Science returned 11’440 documents – there is no single R package for building, exploring, mapping and plotting functional trait spaces. To fill this gap, we propose `funspace`, an R package to handle functional trait space analyses using bivariate or multivariate trait data. Due to the very little difference in using `funspace` with the bivariate or multivariate data input (**Fig. 1**), here we present the package functionalities in the context of PCA-based analyses, and we refer to `funspace` documentation and examples for the case of bivariate trait data. Note that bivariate data can equally be raw trait data as well as pairs of coordinates derived from any ordination method (e.g., NMDS). The package consists of three main interconnected modules (**Fig. 1; Table 1**):

**Fig. 1.**
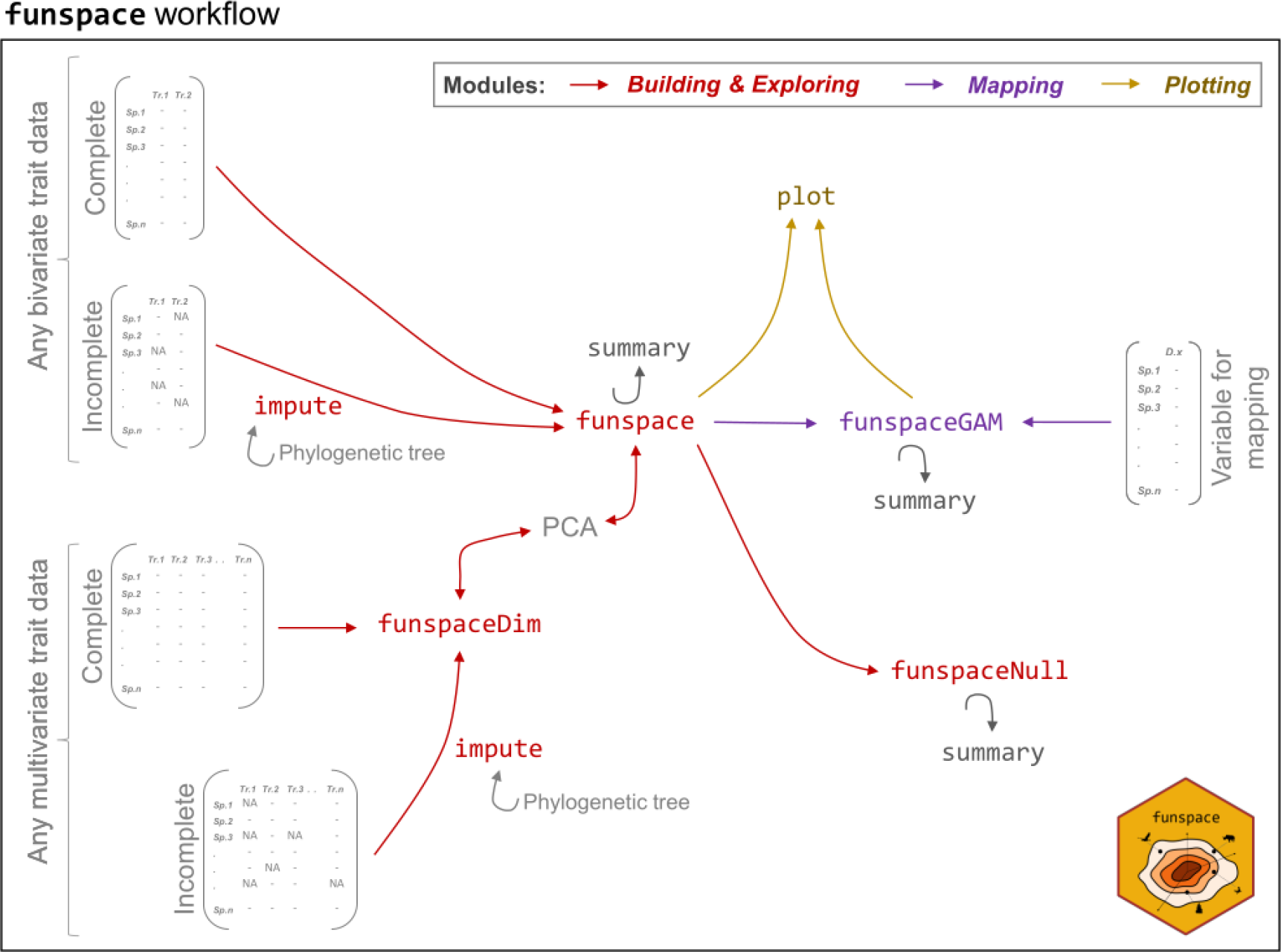
funspace workflow by package module. Functions classification according to package modules is highlighted by color coding. Input data are shown in grey. In case the input trait matrix has missing trait data, the user might want to gap-fill the bivariate or multivariate trait matrix using the `impute()` function, with the option to account for phylogenetic information in the imputation process. Given a multivariate trait matrix, the user needs firstly to assess the number of dimensions that are needed to build a functional trait space using `funspaceDim()`. The second step is running a PCA using `princomp()` before using the other `funspace` functions. The double-headed arrows indicate that the dimensions retained using `fuspaceDim()` are those that should be extracted from the subsequent PCA as well as those used to build the trait space using `funspace()`. In cases where the user decides to identify the trait space dimensions using `funspaceDim()`, and to pass the target PCs to `funspace()` instead of the PCA object, at this point the workflow follows the same path as that of any bivariate trait matrix. That is, the user can directly use a bivariate trait matrix (including coordinates obtained from any ordination method) to run `funspace()`. Because `funspace` is strongly focused in providing graphical representations of functional spaces, in cases when the functional space has higher dimensionalities, two dimensions at a time have to be selected within the `funspace()` function. Once the `funspace()` object has been defined, the user can plot the output (i.e., bivariate functional trait space with kernel density estimates) using the `plot()` function. In case the user wants to test the observed functional trait space against a multivariate normal or a uniform distribution, the `funspace()` object can be passed to `funspaceNull()`. `funspaceGAM()` allows assessing how an additional target variable (e.g., temperature, extinction risk, an additional trait non included in the trait space, etc.) is distributed within the functional trait space previously obtained with `funspace()`. The resulting map of the target variable within the functional space can then be represented graphically as a heatmap using `plot()`. All the modules can handle multiple groups. The package also includes a generic `summary()` function that can be used to print the output of `funspace()`, `funspaceNull()`, and `funspaceGAM()`. Function descriptions are reported in **Table 1**. A full worked example is shown in **Box.1-3**.

**Table 1.** `funspace` functions by module.

| Module | Function | Description |
| --- | --- | --- |
| ***Building & exploring*** | `funspaceDim()` | Identifies the number of dimensions that are needed to build a functional trait space. |
|  | `impute()` | Imputes missing data in a trait matrix using trait-trait relationships and, if specified, phylogenetic information. |
|  | `funspace()` | Builds a functional trait space including multivariate kernel density at different quantiles. Multi-group option available. |
|  | `funspaceNull()` | Tests for statistical difference between the observed functional trait space area vs theoretical areas generated using either a multivariate normal or a uniform distribution. |
| ***Mapping*** | `funspaceGAM()` | Fits GAMs using a target variable as the response variable and functional trait space axes as a two-dimensional smoother predictor. Returns GAM model predictions within a functional trait space. Multi-group option available. |
| ***Plotting*** | `plot()` | If a `funspace()` object is passed as argument, the function prints a functional trait space with multivariate kernel density estimates. If a `funspaceGAM()` object is passed as argument, the function returns a GAM map (i.e., heatmap) of how a target variable distributes within the functional trait space. Multi-group option available. |
| ***Building & exploring AND Mapping*** | `summary()` | Gives a summary of the main features and results of objects created with the `funspace()`, `funspaceNull()`, and `funspaceGAM()` functions. |

Full information on functions' arguments can be found at .....

1. ***Building and exploring module*:** it consists of multiple functions to build the functional trait space from bivariate or multivariate input data using multivariate kernel density methods, and to analyze the main features of a functional trait space (e.g., functional diversity indexes, testing against null models). If specified, all these analyses can be iterated across levels of a grouping variable (e.g., within groups such as families or populations, etc.).
2. ***Mapping module*:** it consists of one function that uses GAMs to statistically test for the link between a target variable and the position of species within the functional trait space. The function handles multiple groups as well.
3. ***Plotting module*:** it consists of a generic plot function that can receive as input the objects built in either of the previous modules. In case the input is the object built in module 1 (building and exploring), the function prints a bivariate (or pairs of dimensions in case of multivariate) functional trait space displaying bivariate kernel density estimates (for single or multiple groups). If the input is an object built in module 2 (mapping module), then the function prints a heatmap depicting how a target variable is distributed within the functional trait space (for single or multiple groups).

### Module 1: Building and exploring a functional trait space

`funspace` includes three functions to build and explore a functional trait space: `funspaceDim()`, `impute()`, and `funspace()` (**Table 1; Fig. 1**). Here we only illustrate the main features of the functions when the input is a PCA object and refer to the `funspace` documentation (available at ….) for full details on the functions’ arguments and input data. However, note that the same exact procedures can be applied to any bivariate trait data (i.e., pairs of raw traits). An example of how to use the functions constituting module 1 is reported in **Box.1**.

`funspaceDim()` allows the user to identify the number of dimensions that are needed to build a functional trait space. The identified dimensions are those that minimize redundancy while maximizing the information contained in the trait data based on the method proposed by Horn (1965) (see Introduction). Briefly, this method contrasts the eigenvalues produced through PCAs run on (30 * (number of variables)) random datasets with the same number of variables and observations as the input dataset. Eigenvalues > 1 are retained in the adjustment. `funspaceDim()` implements the Horn test via the `paran()` function of the R package `paran` (Dinno, 2018). A trait matrix is the only input needed to run the function. `funspaceDim()` returns the number of dimensions to be retained from the subsequent PCA (**Fig. 1**) and prints this information in the R console.

If the input trait matrix (either bivariate or multivariate) contains missing data, the user can fill the gaps using the `impute()` function. By default, `impute()` fills the gaps in the dataset using information on trait-trait relationships using random forest models as implemented in the `missForest` R package (Stekhoven & Bühlmann, 2012). Optionally, a phylogenetic tree (an object of class `phylo`) can be included to improve the imputation procedure by accounting for species phylogenetic relatedness together with the information on trait-trait relationships (Penone et al., 2014). Briefly, a given number of phylogenetic eigenvectors derived from the tree (specified by the argument `nEigen`, default is 10) are added to the original trait matrix, and the resulting dataset is then used in the above-mentioned random forest-based procedure. The `addingSpecies` argument allows the user to add to the phylogeny those species that are only present in the trait matrix. If `TRUE` (default is `FALSE`), the `phytools::add.species.to.genus()` function (Revell, 2012) is used to add species to the root of the genus if there are congeneric species in the tree. Those species that cannot be added to the phylogeny are imputed without taking phylogenetic information into account. Users who want to make use of the multiple options available in `add.species.to.genus()` should modify their phylogenetic tree beforehand. The `impute()` output is a list containing the imputed version of the trait matrix and its non-imputed counterpart.

After having defined the number of dimensions that define the functional trait space, and having ran a PCA, the user can build the functional trait space using `funspace()`, the core function of the package. By default, the function needs only one object to run: either a PCA object (obtained using the `base::princomp()` function), or a data frame including species coordinates along selected PCs (as well as any bivariate trait data) (**Fig. 1**). When a PCA object is the input of `funspace()`, by default the function uses the first two PCs, but the user can plot any pairs of PC by modifying the `PCs` argument. Pairs of principal components should be set based on the output of `funspaceDim()`. With this input, `funspace()` estimates the probability of occurrence of trait combinations within the space defined by pairs of principal components using kernel density estimation with unconstrained bandwidth selectors combining the functionalities available in the R packages ks (Duong, 2007) and TPD (Carmona et al., 2019). The grid size for computing multivariate kernel density estimates is defined by the `n_divisions` argument, that sets the total number of divisions of each dimension or trait; since the resulting functional space is bidimensional, the total number of cells in which the functional trait space is divided is equal to `n_divisions`. Note that increasing `n_divisions` results in longer time for computing the functional trait space (this can be an issue particularly when the `funspaceNull()` is used), but it increases the smoothness of the space edges in subsequent plotting (see Module 3). When the input is a PCA object, `funspace()` uses by default the range of the PC axes to build the grid limits in which estimating the multivariate kernel density estimates, but user-defined trait ranges can be used for calculations as well. The probability threshold used for multivariate kernel density estimates (i.e., the boundaries of the trait probability density function) can be set using the `threshold` argument (default = 0.999 corresponding 99.9% of the trait probability density function being retained). If the dataset includes a categorical variable, the same procedure can be applied to each level of the grouping variable by specifying it in the `group.vec` argument. In this case, by default, `funspace()` constraints group bandwidths for plotting to the global bandwidth, to avoid kernel density estimates of each group to exceed the borders of the functional trait space defined using the whole dataset. However, group-specific bandwidths can be generated by specifying `fixed.bw = FALSE`.

Other than the information that is later used for plotting, and when the input data is a PCA object, `funspace()` also returns a table reporting the loadings of the traits in the provided PCA (eigenvectors multiplied by the square root of eigenvalues) (**Table 2**). In this sense, it is important to note that the ‘loadings’ returned by `base::princomp()` are actually eigenvectors, which do not provide information about the amount of variance contained within each principal component. The squared loadings are used to estimate the proportion of the original variance of each trait that is explained by each selected component (specified in the `PCs` argument) as well as by the two selected components (**Table 2, Box.1**). This information can help understanding how well each trait is represented by the selected functional space. In addition, and independently of the data input (PCA object or any bivariate trait matrix), `funspace()` always returns a table reporting the functional richness and functional divergence of the dataset (Mason et al., 2005) (**Table 3, Box.1**). Functional richness is the area occupied by the functional trait space. Functional divergence quantifies the average distance of species from the center of gravity of the functional trait space. These indexes are calculated using the trait probability density approach (Carmona et al., 2016, 2019) across the whole dataset, and, if specified, per each level of a grouping variable. The user can retrieve all the `funspace()` output(s) using the `summary()` function (**Box.1**).

**Table 2.**
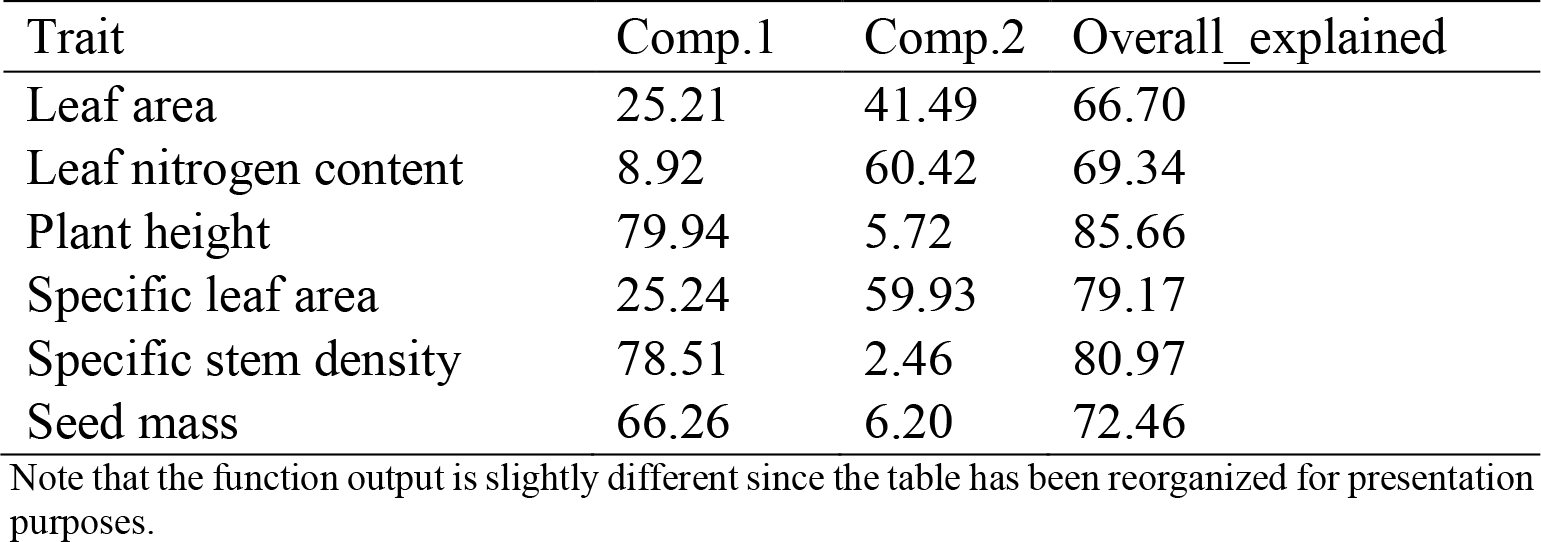
PCA output returned by funspace(). A table summarizing the percentage of variance explained for each trait by each principal component separately (Comp.1 and Comp.2 in this example) and the two components together (Overall_explained). The table is returned as part of the output of `summary(trait_space_global)`, corresponding to the global spectrum of plant form and function using imputed trait information (see Module 1 and **Box.1**). From this output we can see that the first principal component is mainly explaining plant height, specific stem density and seed mass, whereas the second component is more strongly related to leaf traits (leaf area, leaf nitrogen content and specific leaf area). Finally, the `Overall_explained` column reports the percentage of variance explained by the considered space (i.e., the sum of the variance explained by individual components). In this functional space, the quality of the representation of plant height, specific stem density and specific leaf area is better than that of leaf area and leaf nitrogen content.

| Trait | Comp.1 | Comp.2 | Overall_explained |
| --- | --- | --- | --- |
| Leaf area | 25.21 | 41.49 | 66.70 |
| Leaf nitrogen content | 8.92 | 60.42 | 69.34 |
| Plant height | 79.94 | 5.72 | 85.66 |
| Specific leaf area | 25.24 | 59.93 | 79.17 |
| Specific stem density | 78.51 | 2.46 | 80.97 |
| Seed mass | 66.26 | 6.20 | 72.46 |

Note that the function output is slightly different since the table has been reorganized for presentation purposes.

**Table 3.** Functional diversity indexes returned by funspace(). A table summarizing two functional indexes (functional richness and divergence, see Module 1) for the global set of species and for each group contained in the `trait_space_families` example (see Module 1 and **Box.1**). The quantile threshold at which the indexes are calculated is also returned in the output. The table is returned as part of the `summary()` output. In this example we can see that functional richness and functional divergence are larger for the Fabaceae family than for the other families, reflecting the fact that legumes include both herbaceous and tree species (see **Fig. 3**), whereas species within the other families are generally more similar between them.

| Set of species | Threshold | Functional richness | Functional divergence |
| --- | --- | --- | --- |
| Global | 99.90% | 45.55 | 0.56 |
| Fabaceae | 99.90% | 39.92 | 0.58 |
| Lauraceae | 99.90% | 17.13 | 0.46 |
| Pinaceae | 99.90% | 13.19 | 0.39 |
| Poaceae | 99.90% | 28.50 | 0.40 |

Note that the function output is slightly different since the table has been reorganized for presentation purposes.

Finally, the function `funspaceNull()` allows the user to test how the area (i.e., functional richness) of the observed functional trait space differs from that of a null model. Two null models are implemented. (i) The first null model generates data following a bivariate normal distribution (`null.distribution = ‘multnorm’`) with the same mean and variance-covariance structure as the original data. This null model corresponds to the expectation that some trait combinations, specifically those towards the center of the space, are more likely than others, and the resulting space has the approximate shape of an ellipse (Carmona, Bueno, et al., 2021). (ii) Another null model generates a dataset with variables following a uniform distribution (`null.distribution = ‘uniform’`), creating a functional trait space where all trait combinations within the range of the observed functional trait space axes are equally possible, and the space assumes the approximate shape of a rectangle (Díaz et al., 2016). Given a `funspace()` object as input, `funspaceNull()` generates a vector of null model areas across a user-defined number of iterations. Then, using the function `as.randtest` from the `ade4` R package (Dray & Dufour, 2007), it tests for the statistical difference between the observed value of functional richness and the null expectation (considering in both cases the probability threshold value used in the `funspace()` object input). Hypothesis testing options follow the `as.randtest()` specifications. To get meaningful comparisons with the original functional trait space, `funspaceNull()` builds the trait probability density functions at each iteration using the same grid size and bandwidth for computing kernel density estimates as the `funspace()` object. `funspaceNull()` returns: the observed functional richness, the average functional richness of the null model across iterations, and the p-value and standardized effect size associated to the hypothesis test. This information can be retrieved using the `summary()` function. Note that `funspaceNull()` is meant to test the global space against null models, so it does not handle multiple groups.

#### Box.1.

Using `funspace` module 1.

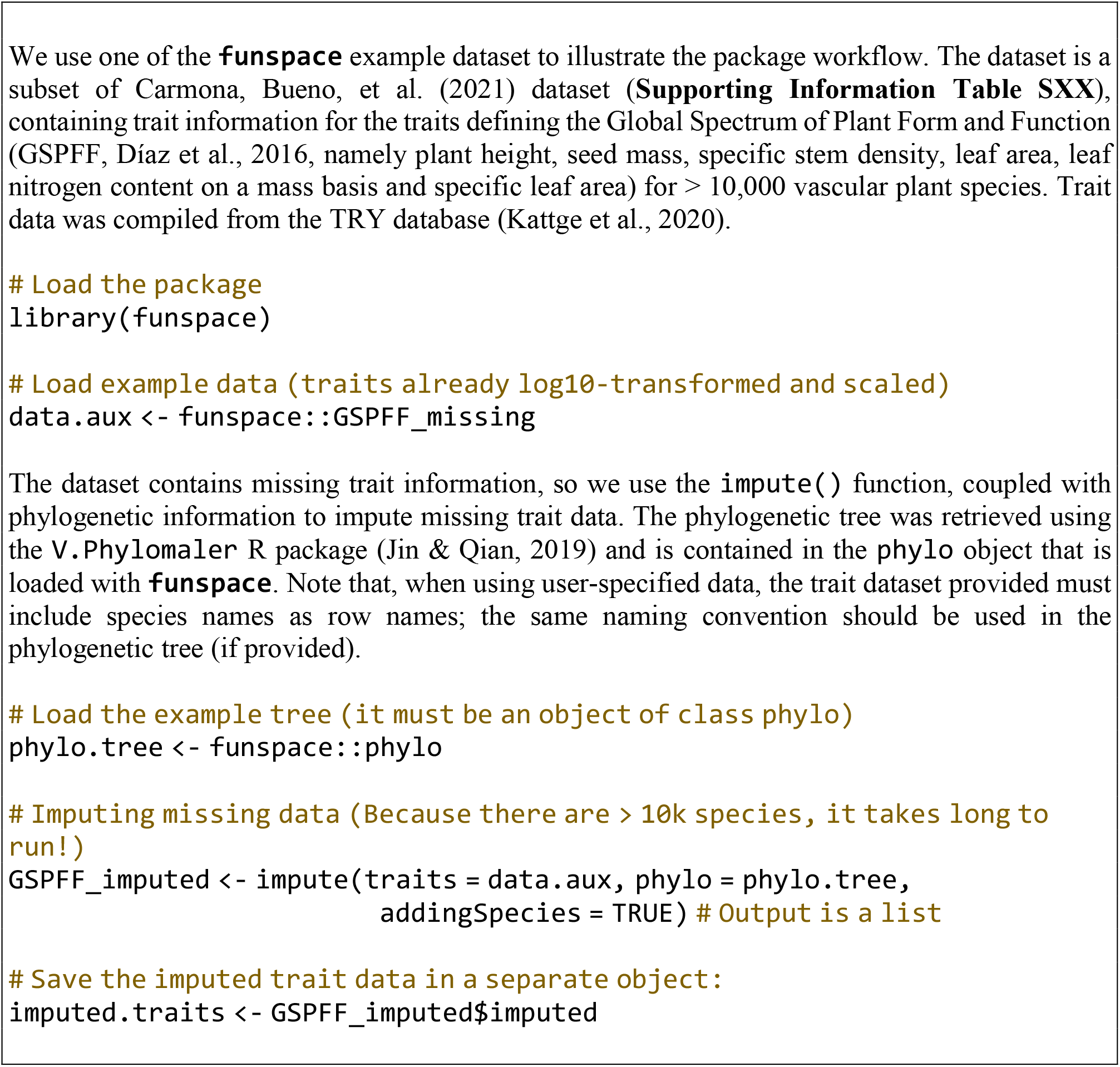
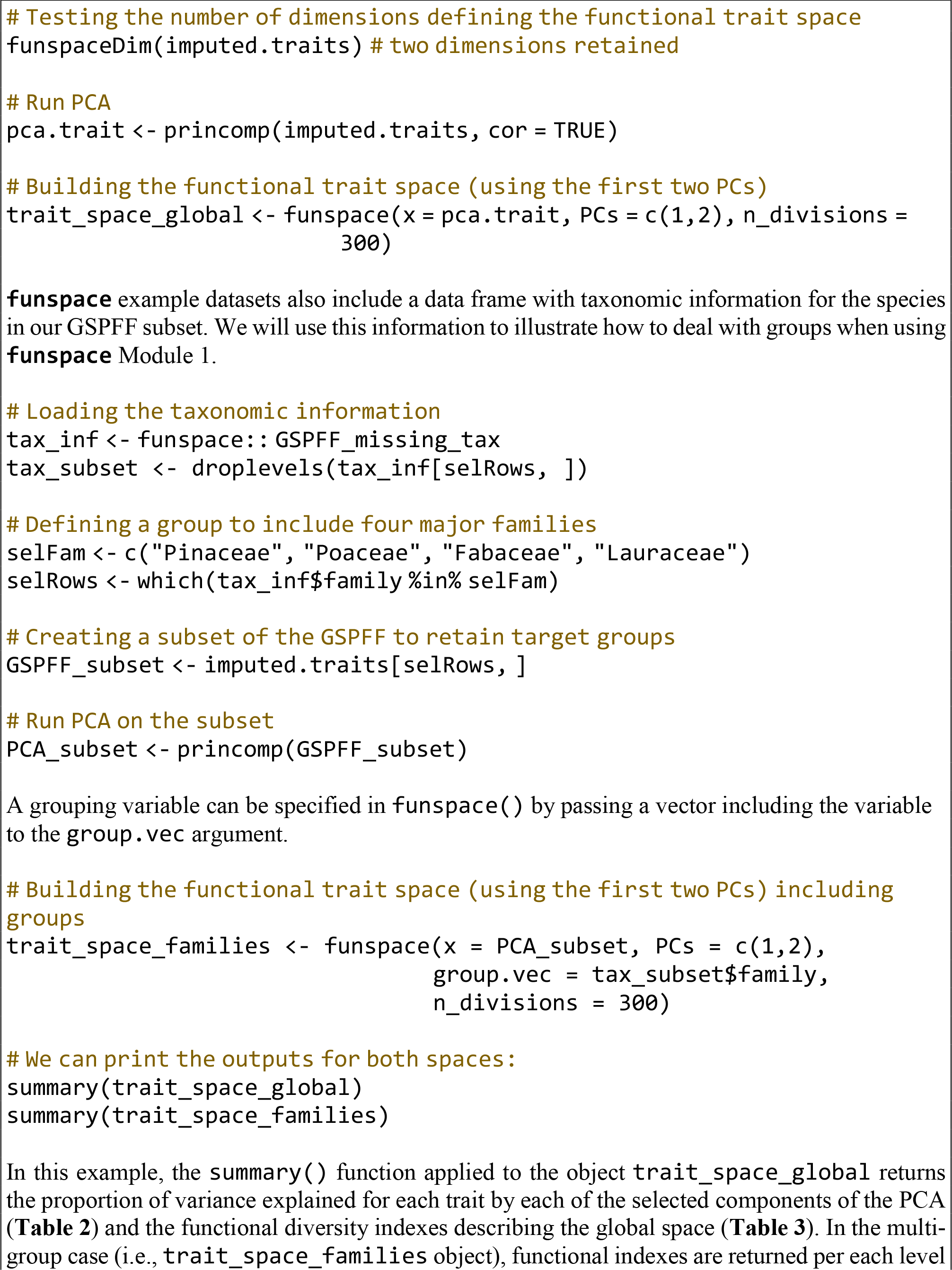
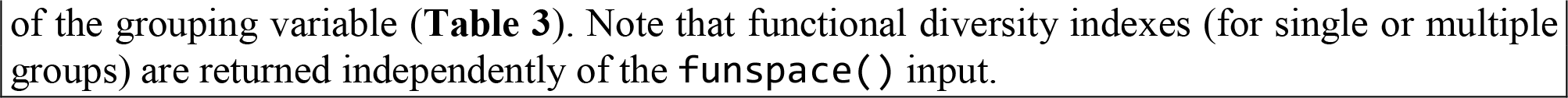

### Module 2: Mapping a functional trait space

`funspace` includes the function `funspaceGAM()` that automatizes GAM modeling steps needed for mapping patterns of a target dimension within the functional trait space (**Table 1, Fig. 1**). An example of how to use `funspaceGAM()` is reported in **Box.2**.

`funspaceGAM()` fits a GAM with a single response variable (defined by the `y` argument) and a two-dimensional smoother defined by the functional trait space axes as the bivariate predictor. GAM specifications are set to the default ones (see `mgcv::gam()` function, Wood, 2017), only the `family` can be specified by the user. Functional trait space axes are inherited by the `funspace()` object specified in the `funspace` argument of `funspaceGAM()`. Given a `funspace()` object, the function automatically defines the functional trait space boundaries and grid size for generating GAM model predictions by inheriting the `threshold` and the grid size information from the `funspace()` object. Note that, the greater is the grid size, the longer is the GAM computational time.

If a grouping variable is specified when running `funspace()` (see Module 1, **Box.1**), GAMs are also fitted per each level of the grouping variable and only within the portion of the space occupied by the data points of each subgroup (see details of `fixed_bw` argument of `funspace()` for different ways of estimating the bandwidth for each group). To avoid fitting models with not enough data, if there are fewer observations in a particular group than specified in the argument `minObs` (defaults to 30), the model is not fitted for that group, and a warning message is returned to inform the user. GAM summary statistics for all data pooled or, if specified, for each level of the specified grouping variable, can be retrieved using the `summary()` function. The output is not shown in the example below because the summary is equivalent to that of the function `mgcv::summary()`.

#### Box. 2.

Using `funspace` mapping module.

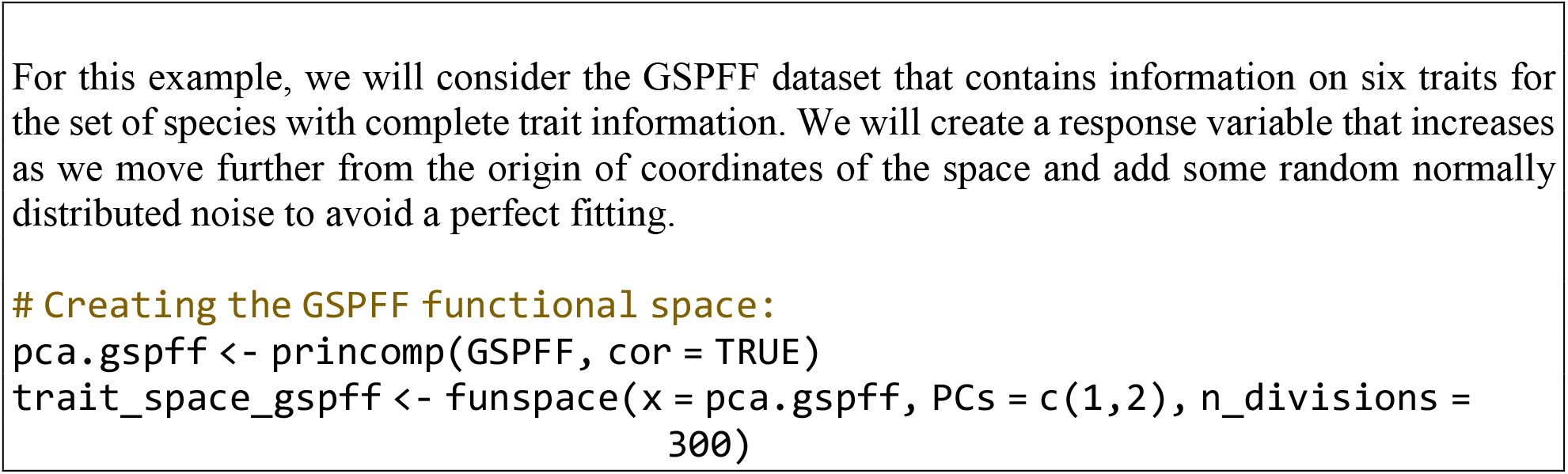
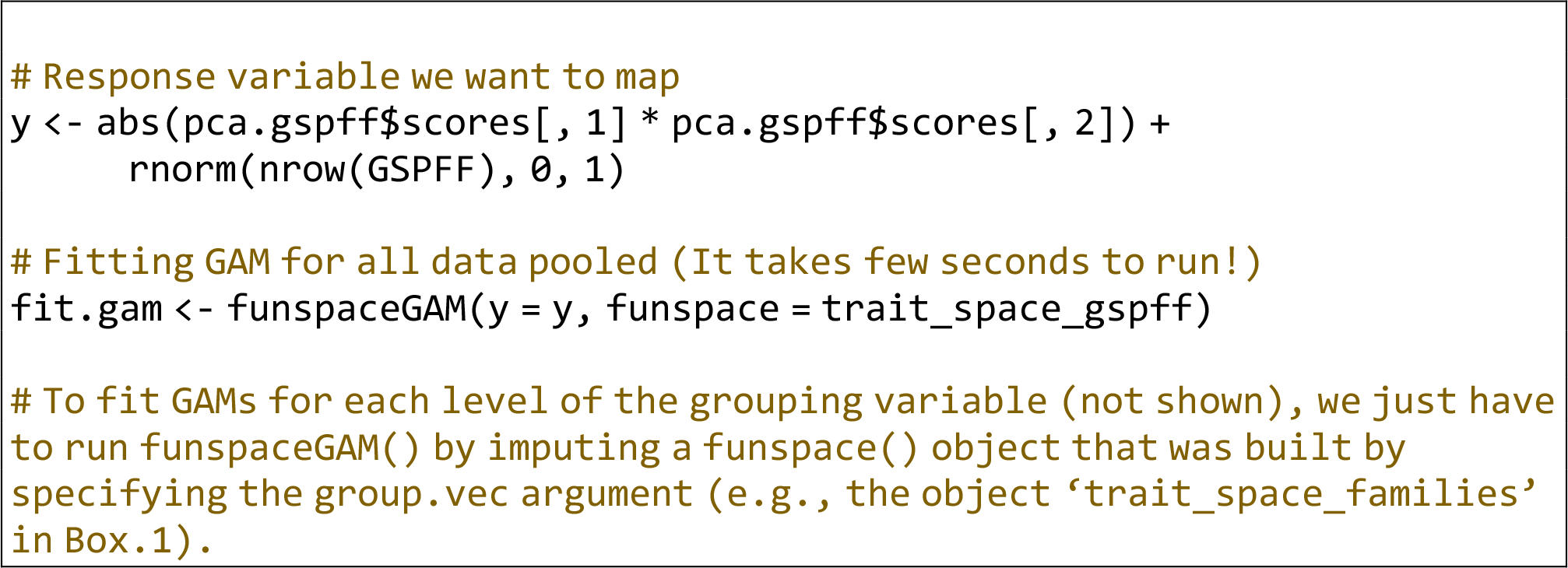

### Module 3: Plotting a functional trait space

`funspace` includes a generic `plot()` function that allows the user to plot objects of the `funspace` class created with the `funspace()` and `funspaceGAM()` functions (see **Box 3** for a worked example).

By default, the only argument needed for plotting is an object of class `funspace` that is assigned to the `x` argument of the `plot()` function. If the plotting input was created with the `funspace()` function, the plotting function returns a functional trait space where colored areas represent the density of occupation of the functional space by the considered species (i.e. the trait probability density function as described in Carmona et al., 2016, 2019; **Fig. 2a**). When the input is the result of the `funspaceGAM()` function, the function prints a heatmap in the functional space depicting the predicted values for the response variable within the boundaries of the trait probability density function (**Fig. 2b**). Note that, in both cases, the resulting plot(s) might appear more or less smooth depending on the number of `n_divisions` set when building the functional trait space using the `funspace()` function (**Box.1**). Larger values of `n_divisions`, will result in smoother plots, but also in higher computational times. An important argument of the `plot()` function is the `type` argument. By default, this argument is set to `‘global’`, and it allows the user to plot the functional trait space for all data pooled (**Fig. 2a,b**). However, if `type` is set to `‘groups’`, and given that a grouping vector has been already specified when building the `funspace()` object (**Box.1**), the `plot()`function creates a plot for each level of the grouping variable in separate panels (**Fig. 3**, see the `fixed_bw` argument of `funspace()`).

**Fig. 2.**
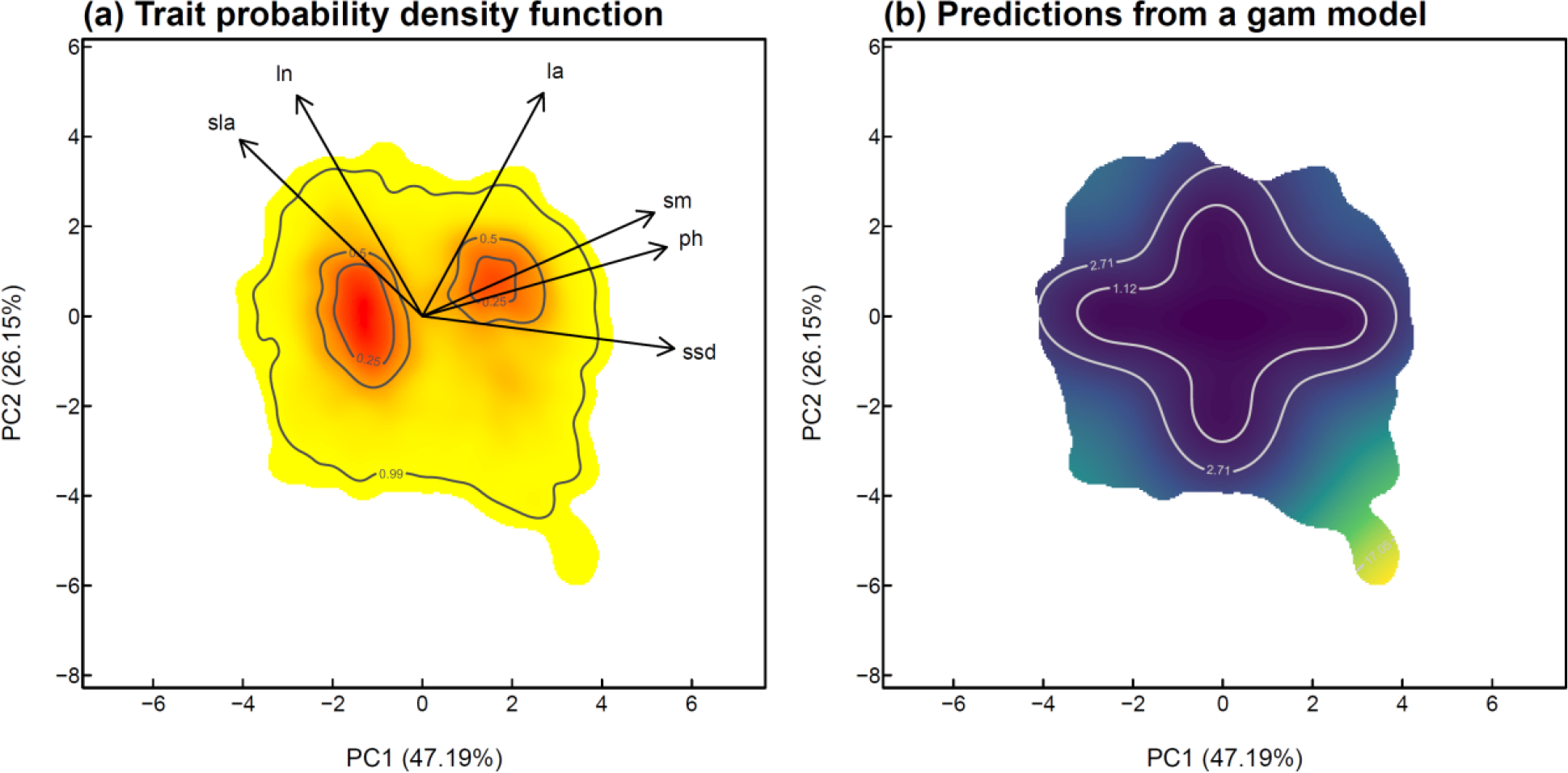
plot() output by input object. **(a)** Plot of a `funspace` object created with the function `funspace()` obtained using all data points in the `GSPFF` dataset. Colors indicate the probabilistic distribution of trait combinations in the functional trait space defined by a PCA (red = high probability; yellow = low probability). Contour lines indicate 0.99, 0. 50, and 0.25 quantiles of the probability distribution. The output shows that there are two hotspots (corresponding, from left to right, to herbaceous plants and angiosperm trees, see Díaz et al., 2016). The variance explained by each component and the loadings of the original traits are also shown. **(b)** Plot showing the predicted values for a variable (see Box 3 for explanations) within the same functional space; the input was created using `funspaceGAM()`. The heatmap shows the predicted values for a variable with increasing values as we move away from the center of coordinates of the functional space. Contour lines are the quantiles of the GAM predictions expressed in the same unit as the response variable. The code to reproduce each panel is available in **Boxes 1-3**.

**Fig. 3.**
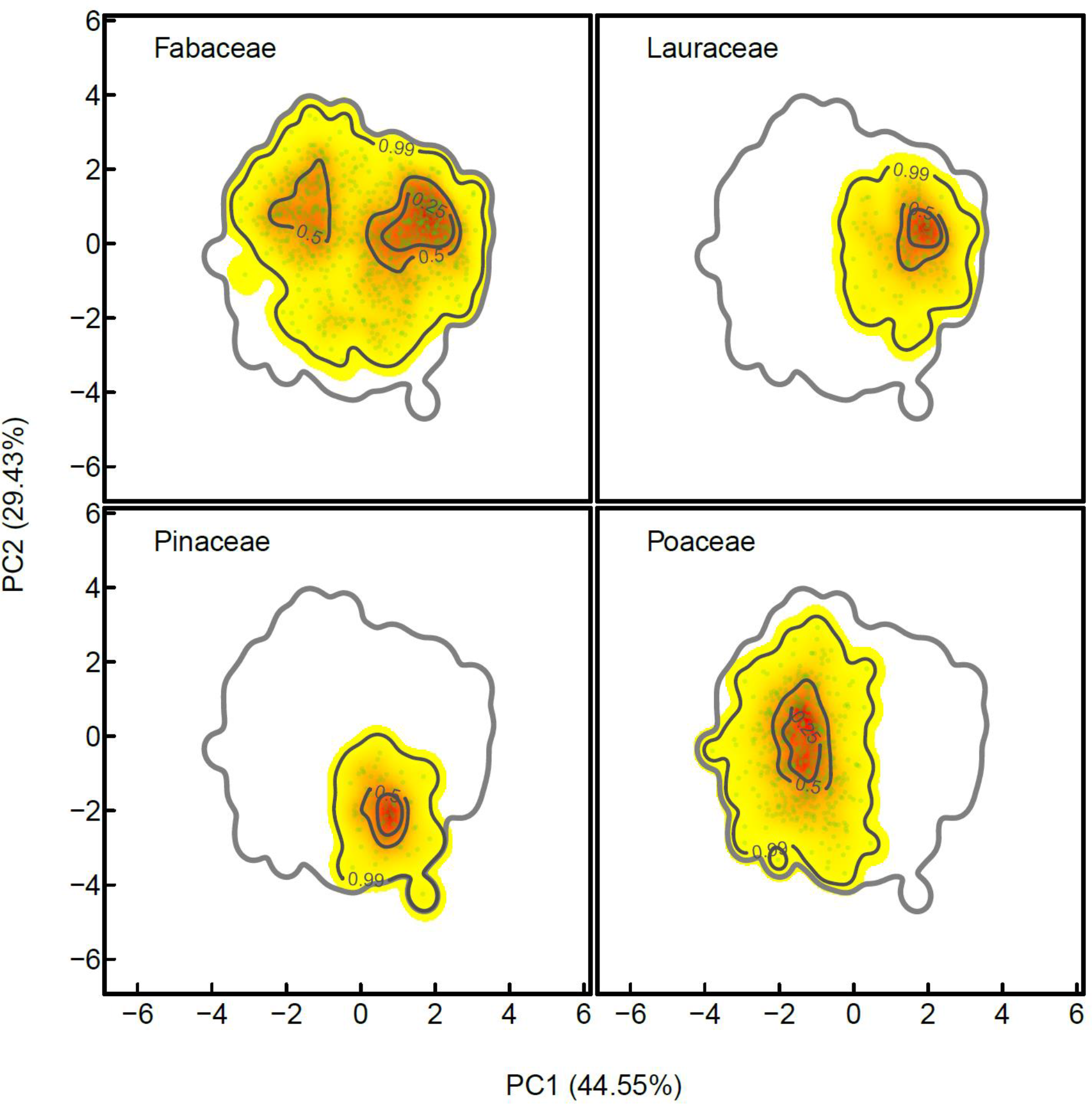
Plot of a funspace object created with the function funspace() including groups for four major plant families (Fabaceae, Lauraceae, Pinaceae, Poaceae). For each family, its corresponding trait probability distribution is represented within the functional space; the 99.9% probability quantile of the global trait probability distribution (i.e. the one including all species from the four families together) is shown to provide a common reference and make comparisons easier. Colors, interpretation of contour lines and explanation of axes is the same as in **Fig. 2a**. The code to reproduce this figure is available in **Box.1** and **Box.3**.

Many graphical parameters can be set for plotting. For example, if the input object comes from `funspace()`, when the argument `quant.plot` is set to `TRUE` (default is `FALSE`), contour lines are drawn at quantiles of the trait probability density function specified by the argument `quant` (by default these are the `threshold` argument that was used in `funspace()` and the 0.50 and 0.25 quantiles). If `quant.plot` is set to `TRUE` when the input comes from the `funspaceGAM()` function, then the contour lines indicate the quantiles of the values of the response variable predicted by the GAM model. When quantile lines are displayed, their features (e.g., type, color) can be set using specific graphical parameters (e.g., `quant.lty`, `quant.col`). Independently of the input object, the gradient palette (with a number of colors specified by the `ncolors` argument, which defaults to 100) can be easily modified by specifying a vector including the two or more colors in the `colors` argument. Plot axes limits can be adjusted by modifying the base plot graphical parameters `xlim` and `ylim`. When the input used to create the `funspace()` object is a PCA, the user can also decide whether to plot arrows representing the trait loadings in the PCA by setting the argument `arrows` to `TRUE` (default is `FALSE`). Full details on plotting options (e.g., points display, etc.) can be found in the `funspace` documentation (….).

#### Box. 3.

Using `funspace` plotting module.

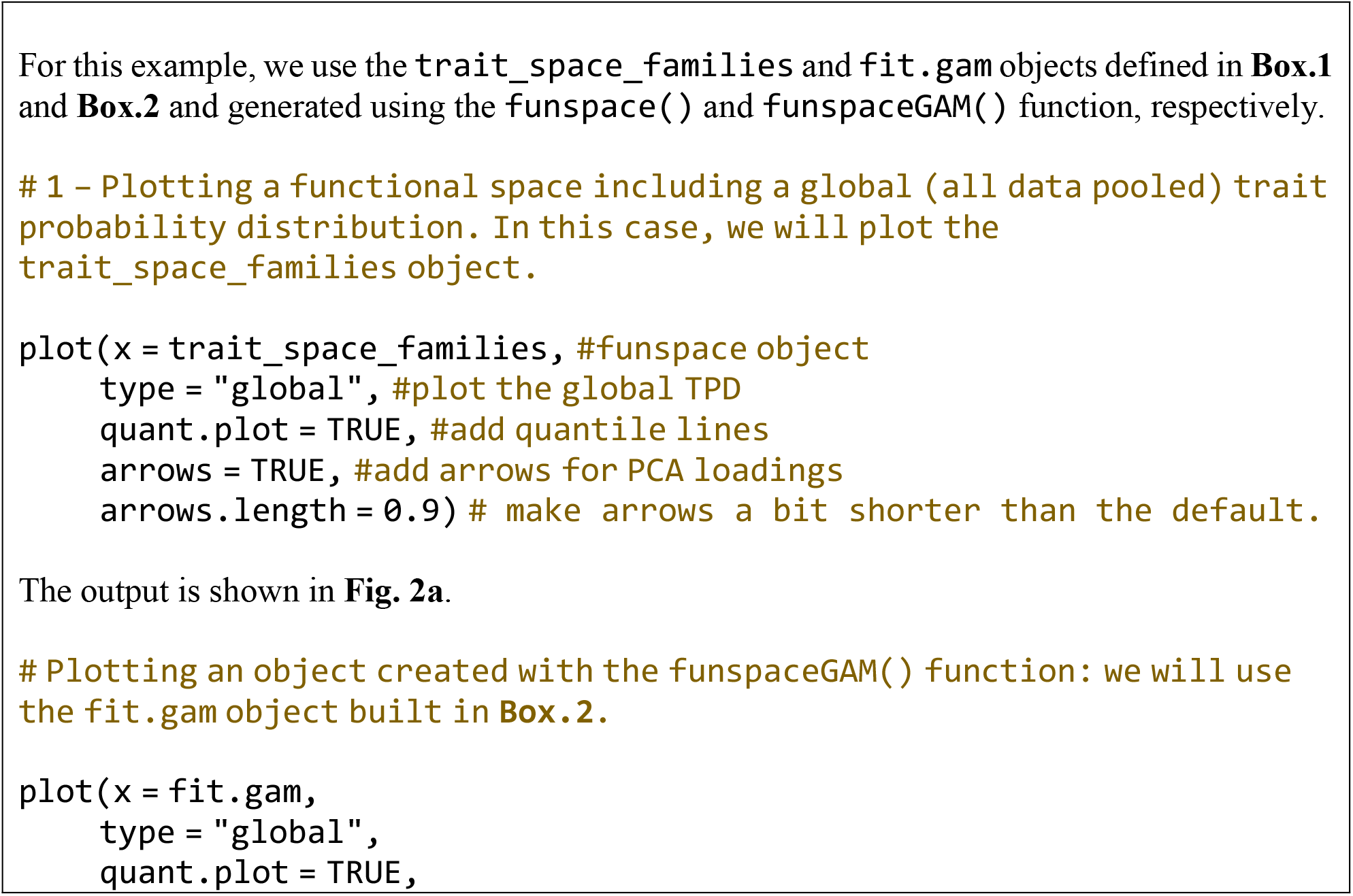
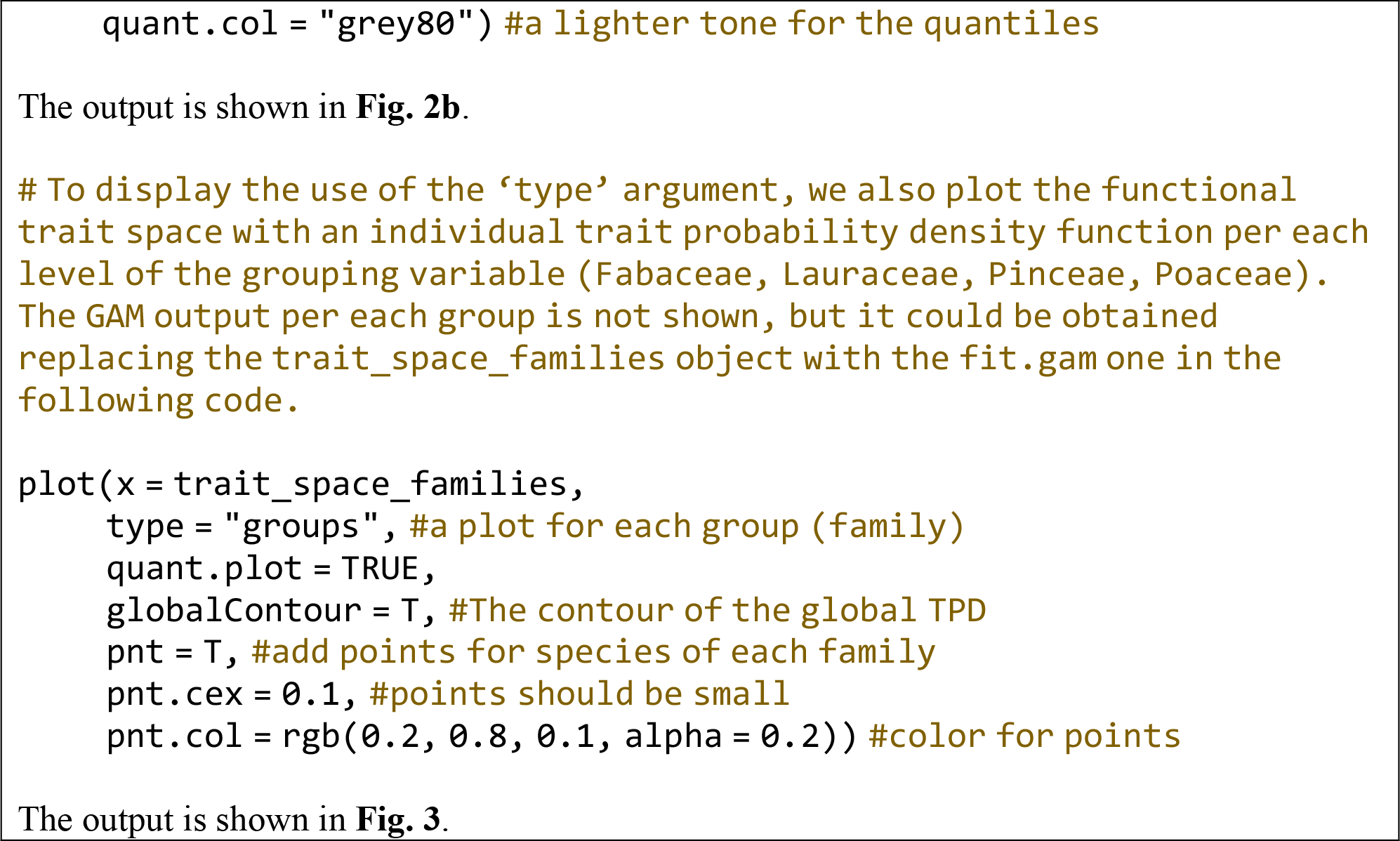

## Conclusions

`funspace` streamlines all the necessary steps to build, explore, map, and plot functional trait spaces, making such analyses readily accessible to any user interested in bivariate or multivariate functional trait analyses. Importantly, `funspace` automatizes most of the procedures needed to run such analyses, and for this reason can be of use to inexperienced R users as well. This package fills an important gap in functional ecological analyses, providing a standardized and reproducible set of procedures that can be used to increase comparability among studies involving functional trait space analyses at different scales and across disciplines. Moreover, this package, due to its easy usage, has the potential to increase the number of studies addressing research questions that require simultaneous consideration of multiple traits and their relationship with multiple ecological variables, such as climate (i.e., functional trait space-environment relationships). Lastly, `funspace` provides key outputs to interpret the main features of any functional trait space and publication-ready plots, increasing the usefulness and potential applicability of this package. We hope that `funspace` will provide the basic tool to build species’ functional traits spaces across the tree of life, to explore the main features of such spaces, and to link them to any biological and non-biological factor that can influence species’ functional diversity.

## Acknowledgments

GP was supported by the grant IJC2020-043331-I funded by MCIN/AEI /10.13039/501100011033, and by the grant PID2021-122214NA-I00 funded by MCIN/AEI/ 10.13039/501100011033 and by FEDER “ESF Investing in your future”, CPC was supported by the Estonian Research Council (PSG293) and the European Regional Development Fund via the Mobilitas Pluss programme (MOBERC40).

## Authors contribution

CPC and GP conceived the idea. All authors equally contributed to the package development. CPC produced the final version of the package. GP and NP wrote the first draft of manuscript. All authors contributed to reach the final version.

## Data availability statement

funspace R package is available at CRAN (https://cran.r-project.org/web/packages/funspace/index.html). All the data that are necessary to run the code reported in the manuscript are available as package example datasets.

